# Identification of Aorta-Specific DNA Methylation Patterns in Cell-Free DNA from Patients with Bicuspid Aortic Valve-Associated Aortopathy and Correlation with Aortic Wall Cell Death

**DOI:** 10.1101/2020.12.20.423686

**Authors:** Ashna Maredia, David Guzzardi, Mohammad Aleinati, Fatima Iqbal, Aiswarya Madhu, Xuemei Wang, Alex J. Barker, Patrick M. McCarthy, Paul W. M. Fedak, Steven C. Greenway

**Author notes:** **Corresponding author:** Dr. Steven C. Greenway, Section of Cardiology, Alberta Children’s Hospital, 28 Oki Drive NW, Calgary, AB, Canada, T3B 6A8. **Conflict of Interest Statement:** We have no conflicts to declare. **Sources of Funding:** University of Calgary and Northwestern University. Greenway is supported by the Alberta Children’s Hospital Research Institute.

## Abstract

**Objective:** In a proof-of-concept study we sought to identify aorta-specific differentially methylated regions (DMRs) detectable in plasma cell-free DNA (cfDNA) obtained from patients with bicuspid aortic valve (BAV)-associated aortopathy.

**Methods:** We used bioinformatics and publicly-available human methylomes to identify aorta-specific DMRs. We used data from 4D-flow cardiac magnetic resonance imaging to identify regions of elevated aortic wall shear stress (WSS) in patients with BAV-associated aortopathy undergoing surgery and correlated WSS regions with aortic tissue cell death assessed using TUNEL staining. Cell-free DNA was isolated from patient plasma and levels of candidate DMRs were correlated with aortic diameter and aortic wall cell death.

**Results:** Aortic wall cell death was not associated with maximal aortic diameter but was significantly associated with elevated WSS. We identified 24 candidate aorta-specific DMRs and selected 4 for further study. A DMR on chromosome 11 showed acceptable specificity for the aorta and correlated significantly with aortic wall cell death. Plasma levels of total and aorta-specific cfDNA did not correlate with aortic diameter.

**Conclusions:** Elevated WSS created by abnormal flow hemodynamics is associated with increased aortic wall cell death which supports the use of aorta-specific cfDNA as a potential tool to identify aortopathy and stratify patient risk.

**Date and Number of Institutional Review Board Approval:** REB17-0207

## INTRODUCTION

The aortopathy that occurs as a consequence of a congenitally bicuspid aortic valve (BAV) is associated with a risk of dissection, aneurysm or rupture.^1^ The etiology of BAV-associated aortopathy appears to be multifactorial and related to inherent genetic defects combined with hemodynamic wall shear stress (WSS) created by turbulence across the abnormal valve.^2-5^ Increased aortic WSS leads to the degradation of elastin and the dysregulation of extracellular matrix proteins which are linked to smooth muscle cell death.^6-8^ With progressive aortopathy, aortic valve replacement surgery is often recommended but patient selection is controversial and often dependent upon individual and institutional practice.^9-13^ A blood-based assay in combination with advanced imaging techniques to identify those who would most benefit from prophylactic surgery would be a significant advance.^14^

Cell-free DNA (cfDNA) refers to fragments of genomic DNA released into the blood during cellular apoptosis.^15-17^ The accessibility of plasma cfDNA and its retention of genetic and epigenetic changes has resulted in the development of cfDNA-based diagnostic assays for diverse human diseases and applications.^18-20^ DNA methylation is an important regulator of gene expression and determinant of cell specialization^21^ and tissue-specific differentially methylated regions (DMRs) have been identified for multiple human cells, tissues and organs.^19^ We hypothesize that aorta-specific DMRs detectable in cfDNA will allow the non-invasive assessment of disease and prediction of important clinical events and enable precision medicine for optimal management of patients with aortopathy and highly variable individual risk.

In this proof-of-concept study we identified novel and unique aorta-specific DMRs that could be measured in human plasma cfDNA obtained from patients with BAV-associated aortopathy. We identified the relationship between elevated WSS and cell death in the ascending aorta of human BAV patients to demonstrate the biological rationale for cfDNA as a biomarker of aortopathy and identified an association between aorta-specific cfDNA levels, WSS and aortic wall cell death.

## MATERIAL AND METHODS

### Patient data

This study was approved by the Research Ethics Boards at the University of Calgary and Northwestern University. After obtaining informed written consent, blood and matched tissue samples from adult patients with BAV undergoing aortic surgery were obtained (Table 1). All patients (n=23) and healthy age-matched controls with tricuspid aortic valves and no documented cardiovascular disease (n=10) underwent 4D-flow cardiac magnetic resonance imaging (CMR) to visualize aortic blood flow patterns, generate WSS heat maps and calculate WSS values as previously described.^8^ Data from the healthy controls (who did not undergo surgery) were used to construct a physiologically normal heat map of aortic flow and regions of depressed, normal and elevated WSS were determined for each BAV patient relative to this control map. Aortic wall tissue samples collected during surgery were flash frozen and then paraffin-embedded.

**TABLE 1.** Patient clinical characteristics.

| Parameter | Study Population |
| --- | --- |
| Age (years) | 49.3 ± 14.9 |
| Male (%) | 20 (87) |
| BAV Classification: |  |
| Type 0 (lateral) | 1 |
| Type 1 (RL) | 15 |
| Type 2 (RN) | 7 |
| Maximum aortic diameter (cm) | 4.67 ± 0.52 |
| Aortic Valve Surgical Procedure: |  |
| Repair | 3 |
| Replacement | 20 |
| Ascending Aorta Surgical Procedure: |  |
| Ascending aorta replacement | 20 |
| Root replacement | 2 |
| Hemi-arch reconstruction | 1 |

Data reported as n or mean with standard deviation. *RL*, right-left coronary cusp fusion; *RN*, right-
non coronary cusp fusion.

### Isolation of cfDNA

Adult BAV patients were recruited and 8-10 mL of blood was collected before the CMR imaging required for their surgery. The blood was centrifuged at 1900 x g for 10 minutes and then the isolated plasma was centrifuged at 4°C at 13,000 RPM for 15 minutes. The resulting plasma was stored at -80°C until use. Patient cfDNA was isolated from 2 mL of plasma using the semi-automated MagNA Pure 24 System (Roche) according to the manufacturer’s instructions. The cfDNA yield was quantified by TapeStation (Agilent) and stored at -80°C.

### TUNEL assay for cell death

Regions of the ascending aorta with the highest and lowest WSS scores were stained for quantification of cell death. Paraffin tissue sections mounted on glass slides were used for terminal deoxynucleotidyl transferase dUTP nick end labeling (TUNEL, Promega) and 4’,6-diamidino-2-phenylindole (DAPI, ThermoFisher Scientific) staining according to the manufacturer’s instructions. Stained tissue slides were imaged using a spinning disk confocal super resolution microscope (SpinSR10, Olympus) at 10X magnification and analyzed using the ImageJ plugin Fiji.^22^ Green (488 nm) TUNEL staining was colocalized to blue (405 nm) DAPI staining and the percent colocalization/cell death was determined.

### Identification of aorta-specific DMRs

Publicly-available methylomes from human hematopoietic cells (n=14), non-aortic tissues (n=21) and aorta (n=1) were obtained from the Roadmap and Blueprint Epigenomic projects.^23, 24^ The software package Metilene identified DNA regions that were significantly differentially methylated (mean difference >10%) based on a Mann-Whitney U test (alpha = 0.05).^25^ Other parameters included a minimum of 4 CpG sites per DMR, <25 base pairs (bp) separating each of the CpG sites and a total DMR length of <100 bp to enable detection in fragments of cfDNA which are ∼140-160 bp in length. A second filtering step identified aortic DMRs with a mean methylation difference >60% for non-aortic tissues and >90% for hematopoietic cells. Comparisons were made separately to enable stringent exclusion of DMRs in peripheral lymphocytes which produce the majority of cfDNA in the circulation. The filtering cutoffs chosen resulted in a sufficient number of DMRs within each interrogated group to enable the identification of DMRs that were common to both groups. Shared DMRs on autosomal chromosomes (sex chromosomes were excluded to avoid dosage effects) identified from the comparison of aorta and non-aorta tissue methylomes (n=446) and from aortic tissue compared to hematopoietic cell methylomes (n=181) provided a total of 23 candidate aorta-specific DMRs (Figure 1).

**FIGURE 1.**
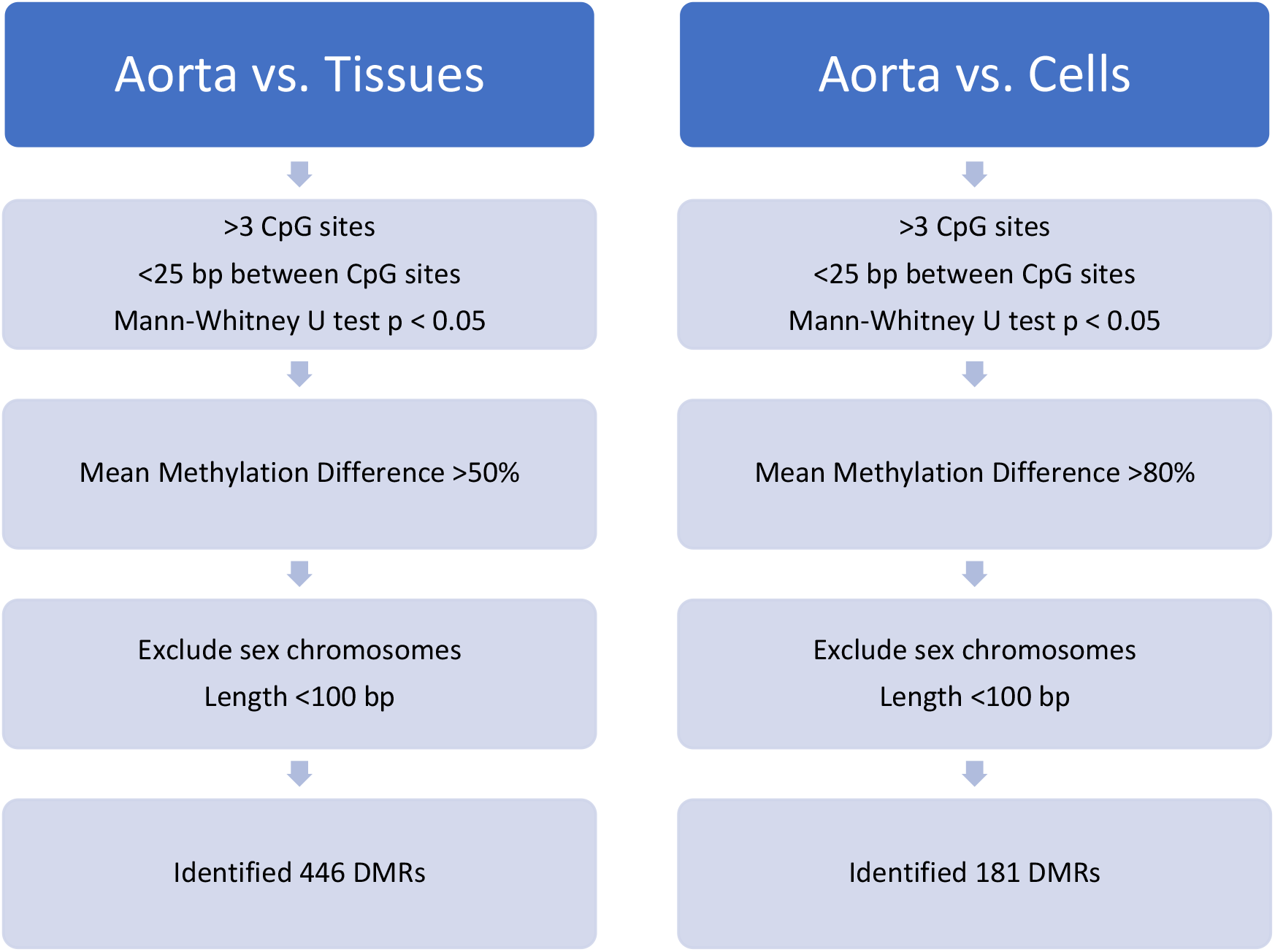
Bioinformatics pipeline used to identify DMRs. Two separate comparisons were conducted in parallel, comparing a human aorta methylome to methylomes from 21 tissues and 14 types of hematopoietic cells to identify 446 and 181 DMRs, respectively. A total of 23 candidate aorta-specific DMRs were shared by both datasets.

For 10 candidiate DMRs with the highest aorta-specificity, sequences containing the DMR of interest and 100 bp upstream and downstream were taken from a methylome of the aorta (GSM983648) generated by the NIH Roadmap Epigenomics project. These sequences were used as input for MethPrimer^26^ to generate primers for amplification of the DMRs from bisulfite-converted genomic DNA (gDNA). Primers did not contain CpG islands and contained at least two non-CpG cytosines. Total product size was constrained to <150 bp to enable amplification from cfDNA. Optimal annealing temperatures were determined for all DMRs. From these, four DMRs, on chromosomes (Chr) 11, 18, 20 and 22 were chosen for further study as these regions amplified readily with no off-target products (Table 2).

**TABLE 2.** Four candidate aorta-specific DMRs.

| <u>DMR</u> |  |  |  | <u>Mean Methylation Difference (%)</u> |  |
| --- | --- | --- | --- | --- | --- |
| Chr | Start Position | Length | CpGs | Non-aortic tissues | Hematopoietic cells |
| 11 | 3,168,734 | 73 | 6 | 70.60 | 92.75 |
| 18 | 74,171,459 | 46 | 8 | 74.26 | 91.27 |
| 20 | 45,860,400 | 65 | 6 | 86.88 | 81.01 |
| 22 | 27,999,645 | 72 | 7 | 75.36 | 91.28 |

Data presented for the four DMRs studied include chromosomal location, length (in base pairs), number of CpG sites and methylation differences between the aorta and pooled non-aortic tissues and hematopoietic cells as determined computationally. *Chr*, chromosome; *CpG*, cytosine-guanine nucleotides.

To confirm the specificity of our DMRs identified *in silico*, we used a commercial panel (Zyagen) of gDNA from human brain, colon, esophagus, small intestine, kidney, liver, lung, pancreas, stomach, skin, spinal cord, skeletal muscle, spleen and thymus to compare methylation levels for all 4 DMRs across tissues in comparison to aorta tissue. Patient cfDNA and panel gDNA was bisulfite converted using the Epitect Bisulfite Kit (Qiagen) with elution performed twice using 20 μL of buffer EB warmed to 56°C to improve yields. Bisulfite-converted DNA was used for PCR amplification of the DMRs and products were visualized on a 3% Tris-Acetate-EDTA agarose gel and quantified by TapeStation (Agilent).

### Next-generation sequencing

For quantification of methylation, PCR amplicons were pooled in equimolar concentrations for sequencing on an Illumina MiSeq. Libraries were sequenced using the MiSeq reagent kit v3 (150 cycles) to produce 2 x 75 bp paired-end reads. The quality of each sequenced pool was assessed using FastQC.^27^ The bisulfite sequencing plugin for the CLC Genomics Workbench (Qiagen) was utilized to analyze the reads which were mapped to the hg19 reference genome with a minimum acceptable alignment of >80%. Methylation levels were determined for each individual CpG site within each DMR and mean methylation levels were calculated.

### Determination of cfDNA concentrations

The concentration of cfDNA (ng/mL) extracted from 2 mL of patient plasma was divided by 0.00303 ng (mass of a single human genome) to determine the total concentration of cfDNA (copies/mL) in recipient plasma.^19^ To determine the absolute concentration of each DMR, the copies/mL was multiplied by the fraction of unmethylated molecules within the given pool of amplicons as determined from the analysis of the bisulfite-sequenced reads.

### Statistics

Statistical analysis was performed using GraphPad Prism 8.0. The percentage of cell death in the aortic tissue is presented as a mean ± standard deviation. The Shapiro–Wilk test was used to determine normality. For cell death, a paired comparison between regions of normal and elevated WSS was conducted for each patient using a Wilcoxon signed rank test and the comparison between pooled patients was performed using the Mann-Whitney test with p <0.05 considered to be significant.

## RESULTS

### Cell death is increased in aortic regions of elevated wall shear stress

For all individual BAV patients, cell death (% TUNEL/DAPI) was increased in regions of the ascending aorta that demonstrated elevated WSS as determined by CMR and this difference was significant for the paired comparisons and pooled data (Figure 2). The mean cell death (mean percent colocalization of TUNEL and DAPI staining) in the regions with normal WSS was 7.98 ± 5.94% compared to 13.74 ± 5.49% in regions with elevated aortic WSS (P = 0.0027). This data demonstrates that regions of abnormal hemodynamics are associated with increased levels of aortic tissue cell death. However, we did not observe a consistent relationship between maximal aortic diameter and WSS or cell death.

**FIGURE 2.**
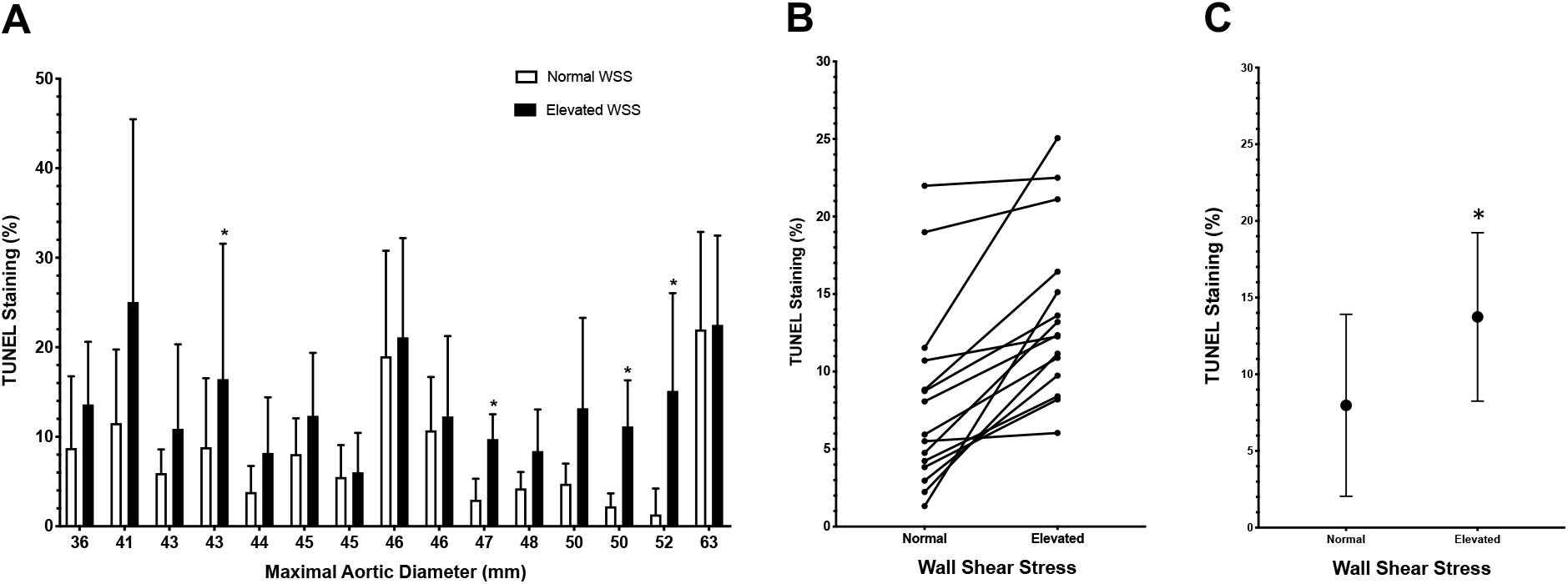
A, Aortic wall cell death (indicated by percentage of TUNEL-positive cells) for 15 patients with varying maximal aortic diameters comparing regions of normal and elevated aortic WSS. Data presented are mean ± SD with * indicating p<0.05 as determined by a Mann-Whitney U test. B, Paired patient data was found to be significantly different (p = 0.00006) using the Wilcoxon signed-rank test. C, Summarized data. Data are mean ± SD with significantly greater cell death in regions of elevated WSS as determined by a Mann-Whitney U test (p = 0.0027).

### Identification of aorta-specific differentially methylated regions

After comparing the aortic methylome to methylomes from 19 tissues we identified 446 putative aortic-specific DMRs with a minimum mean methylation difference of 60%. Comparing the aorta methylome to 14 methylomes from various hematopoietic cells identified 181 aorta-specific DMRs with a minimum mean methylation difference of 90%. There were 24 DMRs on autosomal chromosomes that were common to both datasets and after further testing we selected four DMRs for further study (Table 2). The DMR on Chr 11 (position Chr 11:3,168,734-3,168,832) is 73 bp in length, contains 6 CpG sites and is predicted *in silico* to have the lowest specificity for the aorta in comparison to other tissues but the highest specificity in comparison to hematopoietic cells. The DMR on Chr 18 (position Chr 18:74,171,459-74,171,505) contains 8 CpG sites over 46 bp. The DMR on Chr 20 (position Chr 20:45,860,400-45,860,466) contains 6 CpG sites over 65 bp with the greatest predicted specificity for the aorta in comparison to other tissues. The DMR on Chr 22 (position Chr 22:27,999,645-27,999,717) contains 7 CpG sites over 72 bp. Given the unbiased nature of our enquiry, we were not surprised to see that the regions identified were not specifically associated with genes known to be associated with cardiovascular disease or development. The Chr 11 DMR is found within the *OSBPL5* (oxysterol binding protein-like 5) gene which is an intracellular lipid receptor. The Chr 18 DMR is found within the *ZNF516* (zinc finger protein 516) gene, the Chr 20 DMR is within the *ZMYND8* (zinc finger, MYND-type containing 8) gene and the Chr 22 DMR is not associated with a coding region.

### *In vitro* testing of aorta-specific DMRs

To further test the tissue-specificity of the DMRs that we had identified we obtained gDNA from 14 different human tissues and organs encompassing all three developmental germ layers and compared the measured methylation status in each tissue to the aorta for each DMR. For the Chr 11 DMR, 74.2% of the reads from the aorta were unmethylated, for the Chr 18 DMR 86.8% of the reads were unmethylated, 86.8% of the reads were unmethylated for the Chr 20 DMR and for the Chr 22 DMR, 80.4% of the aorta reads were unmethylated (Figure 3). The Chr 11 and Chr 20 DMRs were mostly unmethylated across all tissues but the Chr 18 DMR was mostly unmethylated in the brain with levels similar to that seen in the aorta and also showed low levels of methylation in the spinal cord, esophagus and colon. For the Chr 22 DMR it was highly unmethylated in skeletal muscle, colon and brain.

**FIGURE 3.**
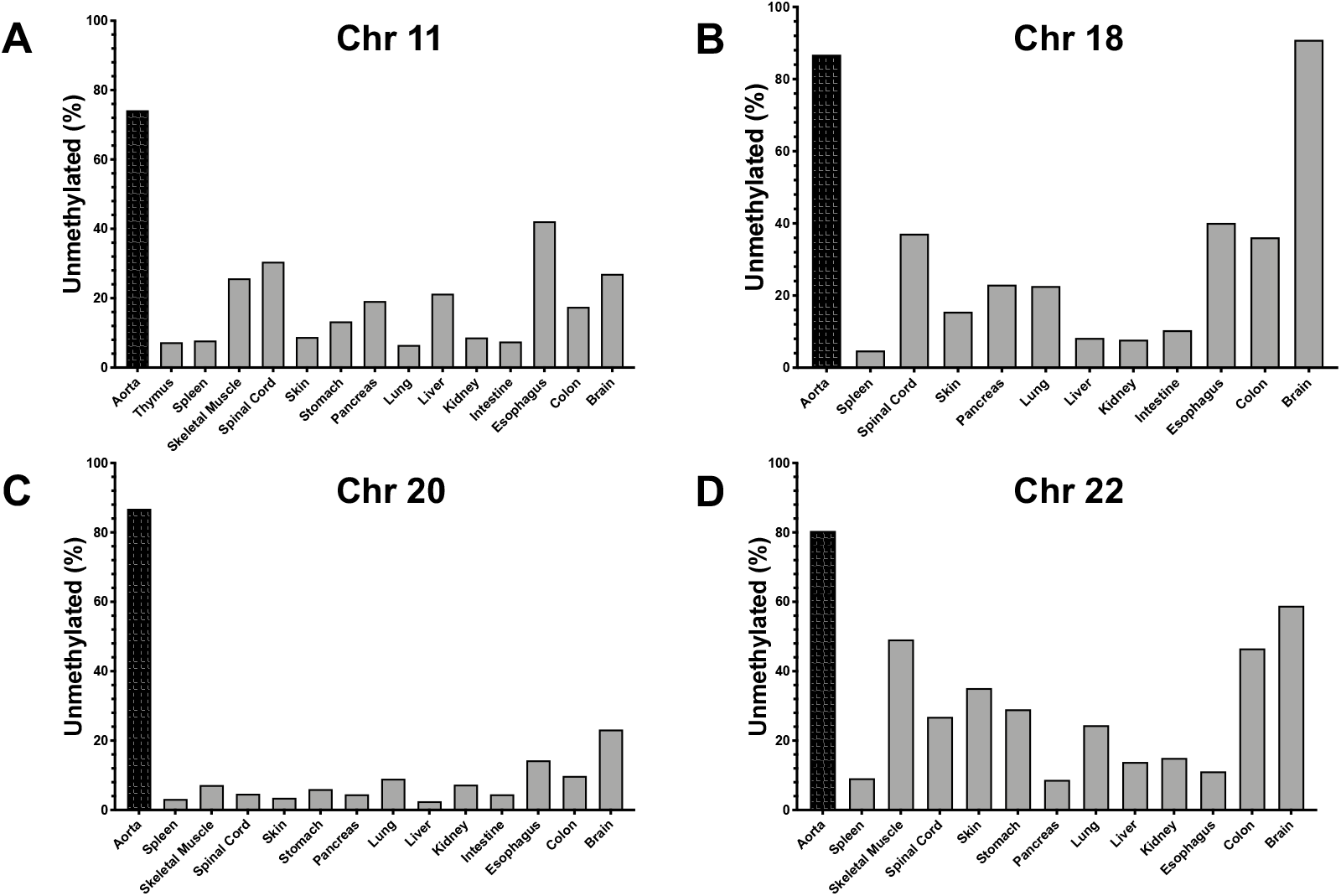
A, Percentage of unmethylated CpG sites for the Chr 11 DMR in aorta and other human tissues and organs. B, Percentage of unmethylated CpG sites for the Chr 18 DMR in aorta and other human tissues and organs. C, Percentage of unmethylated CpG sites for the Chr 20 DMR in aorta and other human tissues and organs. D, Percentage of unmethylated CpG sites for the Chr 22 DMR in aorta and other human tissues and organs.

### Correlation of aorta-specific cfDNA levels with clinical severity of aortopathy

Using cfDNA isolated from patient plasma, the aorta-specific cfDNA levels for 23 BAV patients were determined using the DMRs located on Chr 11, 18, 20 and 22 (Figure 4). None of the aorta-specific DMRs showed an association with aortic diameter as measured using CMR. Similarly, total plasma cfDNA also did not correlate with aortic size. However, levels of aorta-specific cfDNA as determined based on our 4 DMRs did show a positive correlation with levels of cell death in the regions of elevated aortic WSS (Figure 5). This correlation was statistically significant for the Chr 11 DMR (R^2^ = 0.59, p = 0.0035), the Chr 18 DMR (R^2^ = 0.62, p = 0.012) and the Chr 22 DMR (R^2^ = 0.52, p = 0.0078). The Chr 20 DMR did not show a significant correlation (R^2^ = 0.55, p = 0.06). However, our candidate DMRs showed no significant correlation with several histological markers associated with elastin degradation and dysregulation of ECM proteins (concentrations of MMP type 1, 2 and 3, TGFβ-1 and TIMP-1) known to occur in aortopathy.

**FIGURE 4.**
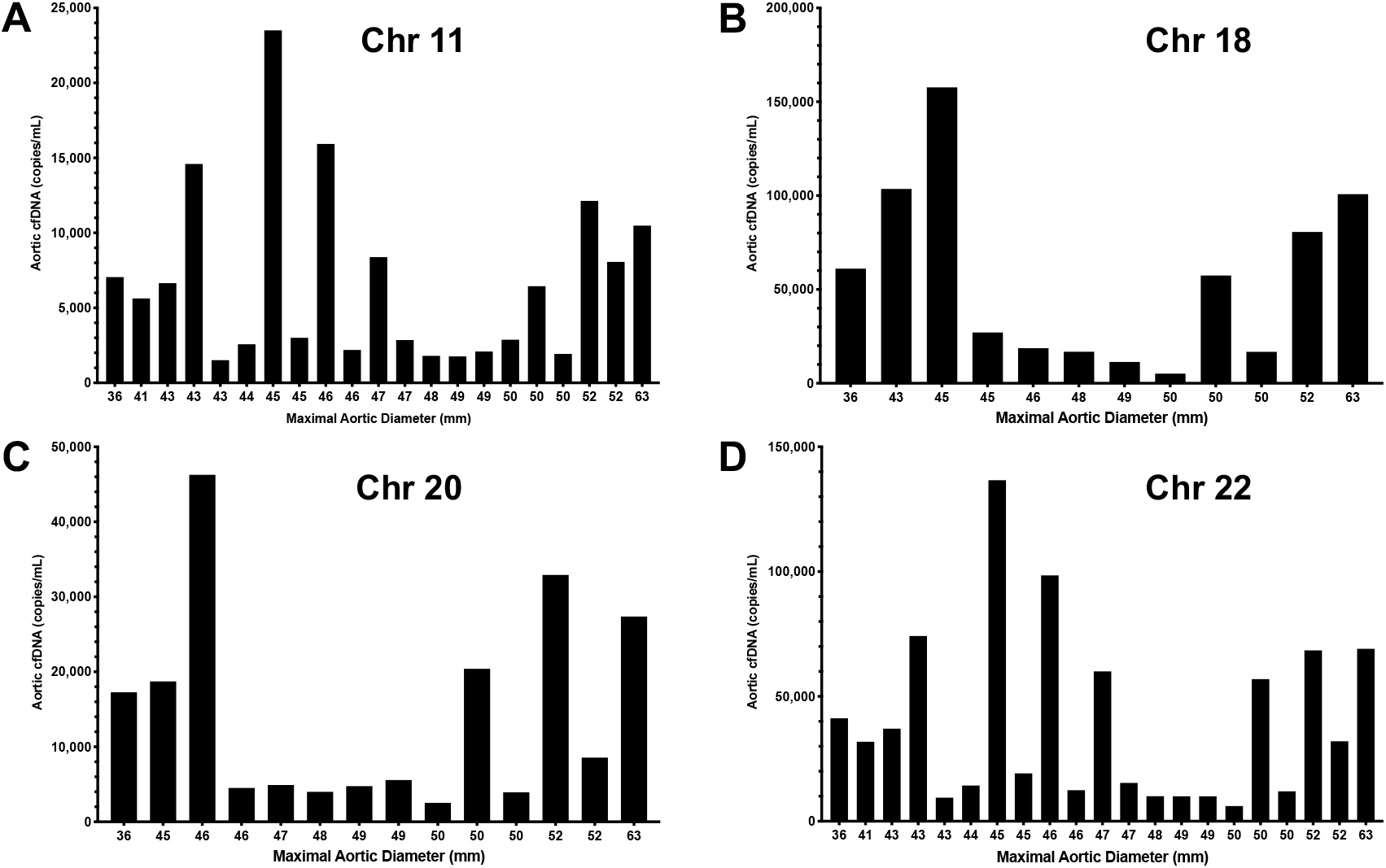
A, Aorta-specific cfDNA concentration as determined by the Chr 11 DMR for 23 patients sorted by maximal aortic diameter. B, Aorta-specific cfDNA concentration as determined by the Chr 18 DMR for 23 patients sorted by maximal aortic diameter. C, Aorta-specific cfDNA concentration as determined by the Chr 20 DMR for 23 patients sorted by maximal aortic diameter. D, Aorta-specific cfDNA concentration as determined by the Chr 22 DMR for 23 patients sorted by maximal aortic diameter.

**FIGURE 5.**
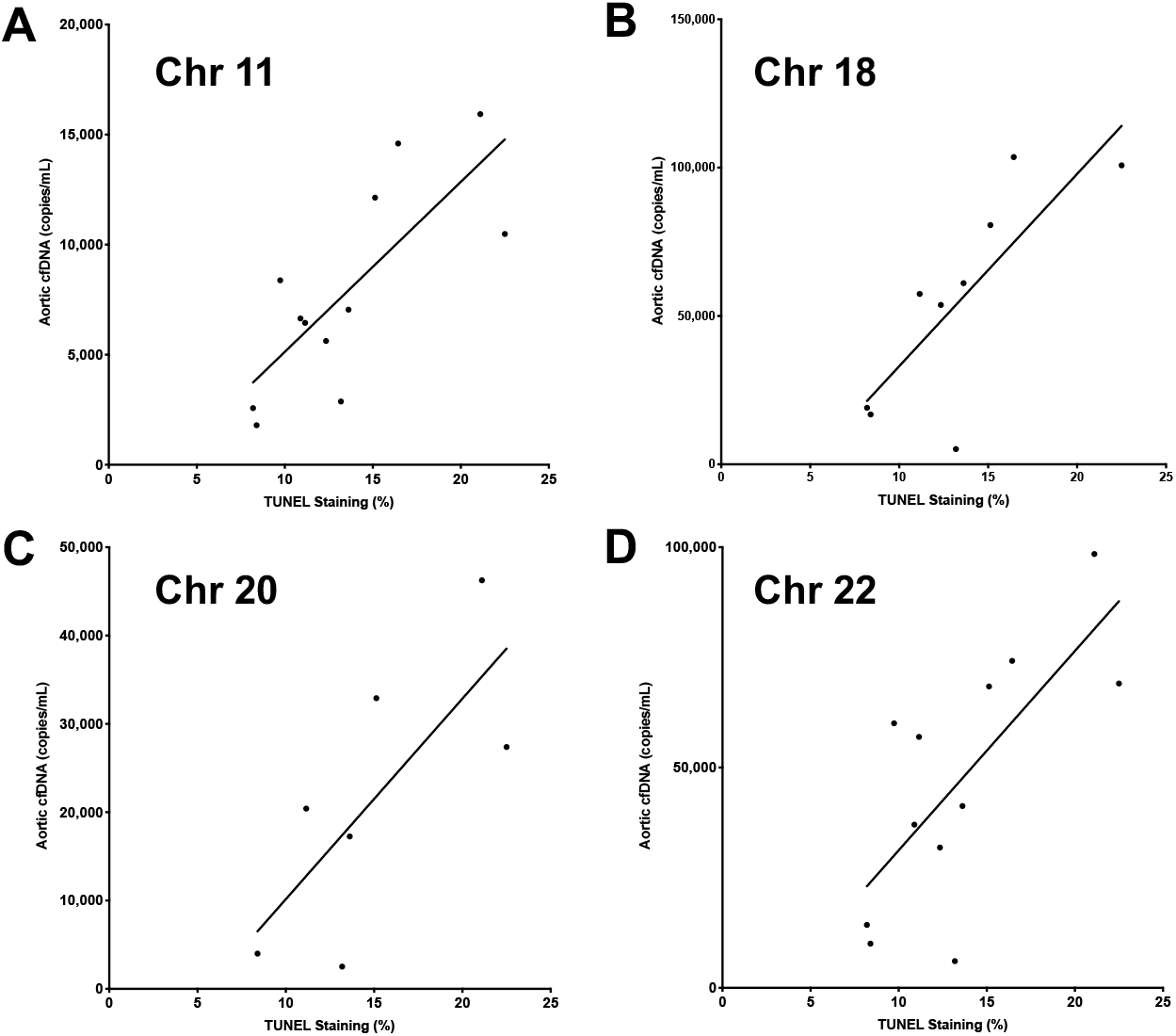
A, Significant correlation (R^2^ 0.59, p = 0.0035) between levels of aorta-specific cfDNA as measured using the Chr 11 DMR and TUNEL staining in regions of elevated WSS. B, Significant correlation (R^2^ 0.62, p = 0.012) between levels of aorta-specific cfDNA as measured using the Chr 18 DMR and TUNEL staining in regions of elevated WSS. C, Non-significant correlation (R^2^ 0.55, p = 0.06) between levels of aorta-specific cfDNA as measured using the Chr 20 DMR and TUNEL staining in regions of elevated WSS. D, Significant correlation (R^2^ 0.52, p = 0.0078) between levels of aorta-specific cfDNA as measured using the Chr 22 DMR and TUNEL staining in regions of elevated WSS.

## DISCUSSION

Using publicly-available human methylomes we bioinformatically identified 24 candidate aorta-specific DMRs. A subset were chosen for further evaluation and the Chr 11 DMR has shown acceptable specificity and significant correlation with aortic wall apoptosis. Our results demonstrate the feasibility of identifying a peripheral blood marker for aortopathy based on DNA methylation patterns and the importance of *in vitro* validation studies.

In this study, the levels of cell death were measured using the TUNEL assay and colocalized to individual cells using nuclear DAPI staining in aortic tissue samples from regions of elevated and normal WSS in BAV patients undergoing surgery. We found that cell death was not associated with maximal aortic diameter but was associated with increased WSS. Our data suggests an association between abnormal hemodynamics and ongoing tissue injury that is independent of aortic size. As these BAV patients experience chronic localized elevated WSS, we hypothesize that the greater levels of cell death in these regions would lead to the depletion of vascular smooth muscle cells and likely negatively impact the integrity of the aorta. Our subsequent data suggests that this increased cell death due to hemodynamic stress is detectable by circulating aorta-specific cfDNA identified through unique DNA methylation patterns.

DNA methylation plays a critical role in regulating gene expression and thus cellular differentiation. Tissue-specific methylation patterns are conserved within a tissue type and to a large degree across individuals.^28^ This consistency is critical for the potential development of a universal, minimally-invasive cfDNA-based assay. From the 24 putative aorta-specific DMRs identified, 4 were selected for detailed study based in part on their computationally predicted specificity for the aorta. The specificity of the 4 DMRs on Chr 11, 18, 20 and 22 was tested *in vitro* using a panel of DNA isolated from multiple human tissues. We found that the methylation differences between the aorta and several tissues and organs were smaller than the *in silico* predictions for many of the DMRs. Importantly, the DMR on Chr 18 was found to be equally hypomethylated in the brain and the aorta and therefore can be eliminated as an aorta-specific biomarker.

Our candidate DMRs were also evaluated using plasma cfDNA obtained from patients with BAV-associated aortopathy undergoing surgery. The levels of both aorta-specific and total plasma cfDNA did not correlate with maximal aortic diameter which is consistent with numerous studies documenting that aortic dimensions (both absolute diameter and rate of progression) are insufficient for assessing the severity of aortopathy and in estimating the underlying risk for aortic rupture.^11, 29-31^ However, although our aorta-specific DMRs did not correlate with aortic dimension, three of them did correlate significantly with levels of cell death in the aorta. This encouraging result provides a rationale for cfDNA as a biomarker for aortic cell death which, as we previously showed, is associated with elevated regions of WSS. Although our candidate DMRs showed no significant correlation with several histological markers associated with elastin degradation and dysregulation of ECM proteins known to occur in aortopathy, the integrity of the aortic wall is impacted by many factors, including apoptosis that leads to the depletion of vascular smooth muscle cells.^32-34^ Regardless of the mechanism of cell death, DNA fragmentation and its release into the circulation is a common endpoint. Thus, the level of aorta-specific cfDNA is potentially an independent and end-stage measure of aortic cell death, regardless of mechanism.

One of the several limitations of our study was the availability of only a single methylome for the aorta. Furthermore, the precise origin of this tissue was unknown. In the future, generating multiple methylomes from the aortic root and ascending aorta may improve the specificity and utility of biomarkers that are developed for BAV-associated aortopathy. Additional limitations were the relatively small, retrospective sample population and the absence of plasma cfDNA from individuals with BAV not undergoing surgery.

In conclusion, elevated WSS created by abnormal flow hemodynamics is associated with increased aortic wall cell death. This finding supports the use of cfDNA as a potential tool to identify aortopathy and stratify patient risk. By leveraging publicly-available data and developing a novel bioinformatics pipeline we identified candidate aorta-specific DMRs that were detectable in cfDNA. Subsequent *in vitro* studies suggest that the Chr 11 DMR possesses acceptable specificity for the aorta and we have shown that it also correlated with the severity of aortic wall apoptosis. However, this intriguing finding needs to replicated and validated in a prospective study.

## Acknowledgements

We would like to thank the nurses and staff involved in the collection of all patient samples and Leo Dimnik for his assistance in the automated isolation of cfDNA. We acknowledge the imaging support provided by the core imaging facility in the Hotchkiss Brain Institute and the sequencing expertise provided by the Centre for Health Genomics and Informatics at the University of Calgary.

## Glossary of Abbreviations

BAV: bicuspid aortic valve
cfDNA: cell-free DNA
Chr: chromosome
CMR: cardiac magnetic resonance imaging
CpG: cytosine-guanine nucleotides
DAPI: 4’,6-diamidino-2-phenylindole
DMR: differentially methylated region
gDNA: genomic DNA
MRI: magnetic resonance imaging
PCR: polymerase chain reaction
TUNEL: terminal deoxynucleotidyl transferase dUTP nick end labeling
WSS: wall shear stress

